# Tissue- and sex-specific small RNAomes reveal sex differences in response to the environment

**DOI:** 10.1101/398784

**Authors:** Alexandra Bezler, Fabian Braukmann, Sean West, Arthur Duplan, Raffaella Conconi, Frédéric Schütz, Pierre Gönczy, Fabio Piano, Kristin Gunsalus, Eric A. Miska, Laurent Keller

**Affiliations:** Department of Ecology and Evolution, University of Lausanne, Switzerland; Swiss Institute for Experimental Cancer Research (ISREC), School of Life Sciences, Swiss Federal Institute of Technology (EPFL), Lausanne, Switzerland; Gurdon Institute, University of Cambridge, Cambridge, UK; Department of Genetics, University of Cambridge, Cambridge, UK; Wellcome Sanger Institute, Wellcome Genome Campus, Cambridge, UK; Center for Genomics & Systems Biology, Department of Biology, New York University, New York, NY; Biologics Discovery California, Bristol-Myers Squibb, Redwood City, CA; Bioinformatics Core Facility; SIB Swiss Institute of Bioinformatics and Centre for Integrative Genomics, University of Lausanne, Switzerland; Center for Genomics & Systems Biology, NYU Abu Dhabi, Abu Dhabi, UAE

**Keywords:** germline sex, *C. elegans*, small RNA, RNA interference, RNA-seq

## Abstract

**Background:** RNA interference (RNAi) related pathways are essential for germline development and fertility in metazoa and can contribute to inter-and trans-generational inheritance. In the nematode *Caenorhabditis elegans* environmental double-stranded RNA provided by feeding can lead to heritable changes in phenotype and gene expression. Notably, transmission efficiency differs between the male and female germline, yet the underlying mechanisms remain elusive.

**Results:** Here we use high-throughput sequencing of dissected gonads to quantify sex-specific endogenous piRNAs, miRNAs and siRNAs in the *C. elegans* germline and the somatic gonad. We identify genes with exceptionally high levels of 22G RNAs that are associated with low mRNA expression, a signature compatible with silencing. We further demonstrate that contrary to the hermaphrodite germline, the male germline, but not male soma, is resistant to environmental RNAi triggers provided by feeding. This sex-difference in silencing efficacy is associated with lower levels of gonadal RNAi amplification products. Moreover, this tissue-and sex-specific RNAi resistance is regulated by the germline, since mutant males with a feminized germline are RNAi sensitive.

**Conclusion:** This study provides important sex-and tissue-specific expression data of miRNA, piRNA and siRNA as well as mechanistic insights into sex-differences of gene regulation in response to environmental cues.

## BACKGROUND

The environment can induce changes in phenotype and gene expression that persist across multiple generations [reviewed in 1]. Such intra-and trans-generational inheritance can pass both through the male and female germlines. Several studies have revealed sex-differences in transmission efficiency of heritable phenotypic changes, yet the underlying molecular mechanisms remain unknown [2–5].

In the nematode *Caenorhabditis elegans*, environmental cues such as starvation, viral RNA or environmental RNA can trigger heritable phenotypic changes that are transmitted via small RNA [6–8]. The mechanisms of trans-generational inheritance are best understood from studies of environmental RNA [5,6,9–11]. In *C. elegans* phenotypic changes induced by environmental RNAi require a double stranded RNA (dsRNA) entering the animal via a dedicated dsRNA transporter [12,13]. Thereafter, exogenous dsRNAs are processed by the conserved nuclease Dicer into ∼22 nucleotide (nt) primary small interfering RNAs (siRNA) and incorporated into Argonaute proteins to form the RNA-induced silencing complex (RISC) [14,15]. This protein-RNA complex binds complementary mRNA sequences and initiates the production of secondary siRNA by RNA-dependent RNA polymerases (RdRP) (Fig 1A) [16–18]. Such secondary siRNAs are 22 nt with a 5’ guanine, thus named 22G RNAs [19]. Primary and secondary siRNA trigger systemic gene silencing of complementary sequences through destabilization of mRNA or translational repression (Fig 1A)[20–22]. Once established, phenotypic changes induced by exogenous dsRNA can be transmitted over multiple generations [9,11,22].

**Figure 1:**
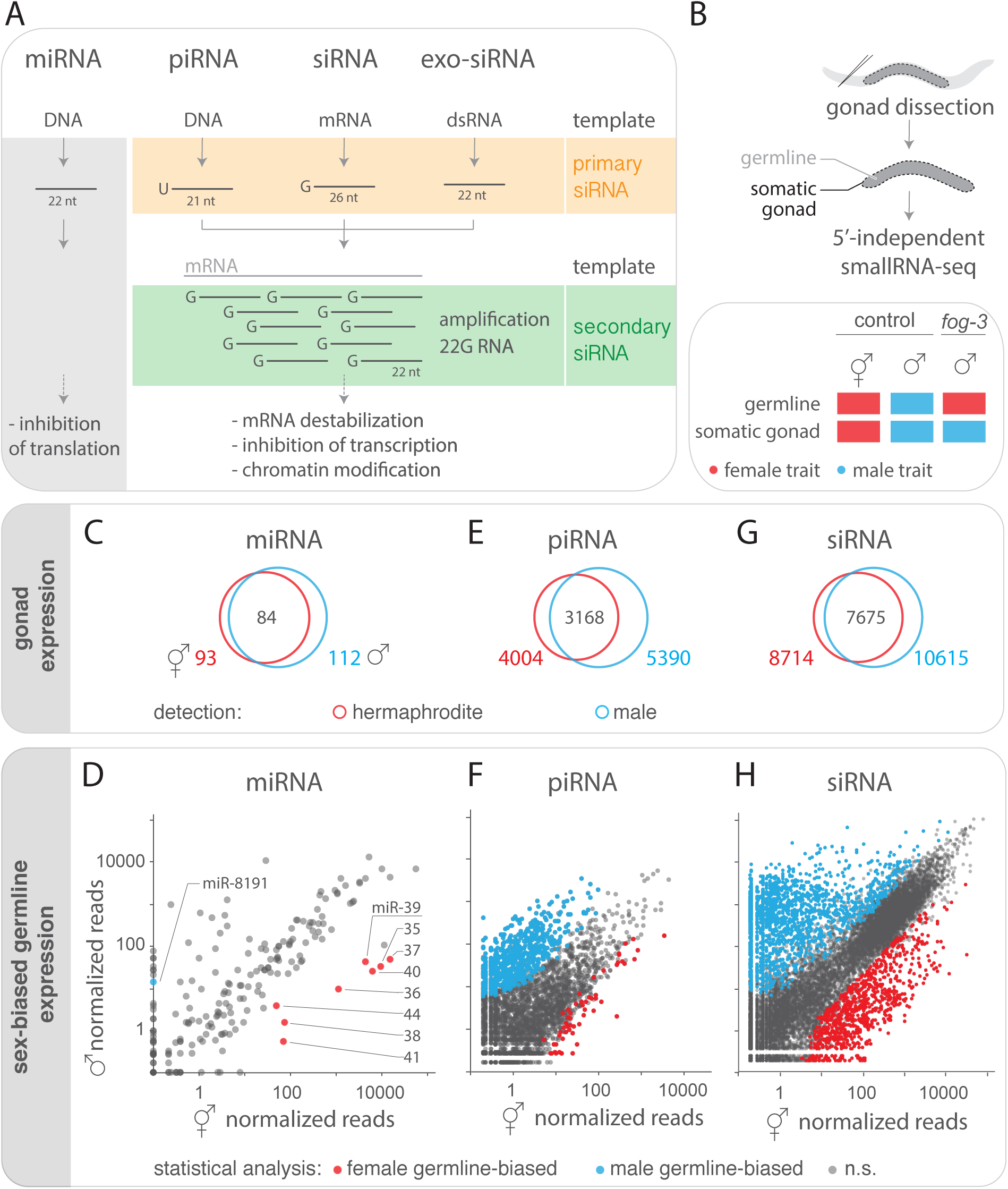
Tissue- and sex-specific small RNA profiling. A) Small RNA types in *C. elegans* categorized by template (DNA or RNA), length and first nucleotide (nt). Distinct primary small RNA types (orange) induce the amplification of secondary 22G RNAs (green) by RNA-dependent RNA polymerase (RdRP) using mRNA as a template. B) Small RNAs were sequenced from dissected adult *C. elegans* gonads containing germline and somatic gonad tissue. Control hermaphrodites and males have sex-specific germline and somatic gonad tissue (red: female traits; blue: male traits). Mutant *fog-3* males have a ‘female’ germline (red) and a male somatic gonad (blue). C,E,G) Gonad expression of miRNA (C), piRNA (E) and endogenous siRNA (G, >5 mean reads) in control hermaphrodites (red circle, n=6 replicates) and males (blue circle n=3). Total number of gonadal small RNAs in each sex is shown as well as the overlap. D,F,H) Sex-differences in expression of mean normalized miRNA (D), piRNA (F) and endogenous siRNA (H) reads in hermaphrodite (n=6 replicates) and male gonads (n=3). (D,H) miRNA and siRNA expression-differences with four-fold difference in abundance that were statistically different (Wilcoxon rank sum test with continuity correction; p adjusted<0.01) between hermaphrodites and males as well as between control males and *fog-3* males (n=2) are highlighted in red (female germline-biased) and blue (male germline-biased). (F) Since piRNA are germline-specific, expression-differences between hermaphrodites and males (n=2) are highlighted in red (female germline-biased) and blue (male germline-biased) without comparison to *fog-3*. See additional file 1 Fig S1 for sex-differences in somatic gonad expression.

On top of their role in trans-generational inheritance, some endogenous small RNA types – such as miRNA, piRNA and siRNA – are essential for development and fertility in both sexes of *C. elegans*. Albeit distinguishable, those small RNA types share some biogenesis and silencing mechanisms (Fig 1A). One major class of small RNA, miRNAs, regulate translation of mRNA targets via partially complementary base-pairing. Many miRNAs are expressed in a tissue-and sex-specific specific manner reflecting functional sex-differences [23–26]. For instance, miR-35 family activity in the female germline is important for progeny viability and fecundity [27–29]. Furthermore, distinct Argonaute proteins associate with additional types of endogenous siRNAs in the male and female germline, suggesting functional sex-differences [19,30–34]. Importantly, germline-expressed PIWI-bound small RNA (piRNA) populations target mRNA and thus maintain genome stability in hermaphrodites and males [35–38]. Such piRNAs come in two flavours: type I piRNA are expressed from two genomic loci have a conserved upstream motif, whereas type II piRNAs lack an apparent upstream motif and are dispersed throughout the genome [39]. In *C. elegans*, another group of endogenous small RNAs are expressed in the gonads of both sexes. Such siRNAs can be distinguished by length and are either (i) primary products of the RNAse III enzyme Dicer/DCR-1 (e.g., 26G RNA) or (ii) secondary products of RNA-dependent RNA polymerases (e.g., 22G RNA) (Fig 1A). Both types of endogenous siRNAs modulate gene expression and are essential for fertility, but very little is known about potential sex-differences in their gonadal expression [30,32,33].

The reproductive tissue of *C. elegans* consists of the germline surrounded by the somatic gonad, and development of both tissues is coordinated by multiple mechanisms [40]. Notably, miRNA activity in the somatic gonad is essential for gonad development and germline proliferation, and thus fertility [41,42]. However, a comprehensive tissue-and sex-specific expression study of gonadal siRNA is lacking.

Males and females also present germline sex-specific differences in response to environmental cues. First, RNAi in hermaphrodite gonads induces very strong knockdown phenotypes [2,43–45], in contrast to anecdotal evidence for mostly no detectable RNAi phenotypes in *C. elegans* sperm [2,43,46–48]. Second, siRNA induced trans-generational silencing is often more efficient through the female than the male germline [2–5]. The underlying mechanisms causing these sex-differences are not well understood.

Here we provide a comprehensive study of both the genetically-determined small RNAome and environmentally-induced siRNA silencing. First, we quantify sex-biased expression of miRNA, piRNA and siRNA in isolated male and female gonads. We further ascribe small RNA sex-differences to the germline or somatic gonad by comparing gonads of each sex to mutant male gonads (*fog-3*) with a female germline. Quantitative analysis of gonadal expression of mRNA and 22G RNA identified genes with low mRNA expression and high 22G RNA levels, a signature compatible with silencing. Second, using environmental RNAi targeting a GFP-sensor we show that germline RNAi silencing efficacy is determined by germline sex. This tissue-specific sex-difference in silencing efficacy was associated with lower levels of RNAi amplification products in male than female gonads. These data provide a mechanistic basis for sex-differences in germline RNAi efficacy with implications for trans-generational inheritance.

## RESULTS

### The small RNAome of male and hermaphrodite gonadal tissues

To identify small RNAs expressed in hermaphrodite and male gonads, we quantified small RNA populations (Fig 1A) from dissected *C. elegans* gonads by high-throughput sequencing (Fig 1B). This approach allowed us to simultaneously analyse and compare tissue-specific expression of miRNA, piRNA and siRNAs of *C. elegans* gonads.

### miRNAs with sex-biased germline or somatic gonad expression

Of the 257 annotated miRNAs in the *C. elegans* genome (assembly WS235, miRBase release 21) we detected 93 in hermaphrodite gonads and 112 miRNAs in male gonads (Fig 1C; minimum 5 mean sense reads across replicates, additional file 2). To uncover quantitative differences in miRNA expression between sexes, we compared normalized read counts between hermaphrodite and male gonads. By using a cut-off of four-fold difference in abundance and a false-discovery rate of 1%, we identified 37 miRNAs with sex-biased gonad expression (Fig 1C).

Since the gonads consist of two tissue types, the germline and the somatic gonad, the observed difference between hermaphrodite and male may stem from expression differences in either tissue. To determine the contribution of the germline, we made use of mutant males that have a feminized germline and a male somatic gonad (i.e., the loss of function allele *fog-3(q849[E126K]* called *fog-3* or ‘feminized male’ for clarity) [49,50]. The comparison of the control male and feminized male gonads allows one to uncover differences that stem only from the germline, since both types of individuals have identical male somatic gonads (Fig 1B). Of the 37 miRNAs with sex-biased gonad expression, nine were differentially expressed between the male and feminized male gonads (Fig 1D). Eight of them (including all form the miR-35-41 cluster and miR-44) were expressed more highly in the feminized germline (*fog-3* gonads) whereas only one (miR-8191) was more highly expressed in the male germline (male gonads) (Fig 1D). This result shows for the first time sex-specific expression of miRNAs in the *C. elegans* germline.

The comparison of hermaphrodite and feminized males permits to also identify the contribution of the somatic gonad to differences in sex expression between hermaphrodite and male gonads, since germlines are identical (both female). This comparison revealed that 16 of the 37 miRNAs with sex-biased gonad expression were differentially expressed between the hermaphrodite and feminized gonads. One miRNA was more highly expressed in hermaphrodites, and 15 miRNAs were more highly expressed in feminized males, indicating sex-differences that stem from the somatic gonad (see additional file 1 Fig S1).

### piRNAs with sex-biased expression

Next, we compared hermaphrodite and male gonads to identify type I and II piRNAs with sex-biased expression. Our cloning technique allowed us to capture (5’-monophosphate-) piRNAs, yet at lower frequency than other types of (5’-triphosphate-) siRNA. We detected sense reads of 4004 annotated piRNAs [36,51] in hermaphrodite gonads and 5390 in male gonads, including 3168 piRNAs present in both sexes (Fig 1E, additional file 2). To uncover putative quantitative differences in piRNA expression between sexes, we compared normalized read counts between hermaphrodite and male gonads. Using a cutoff of four-fold difference in abundance and a false-discovery rate of 1% revealed 66 piRNAs with hermaphrodite-biased expression and 919 piRNAs with male-biased expression (Fig 1F). piRNAs are germline specific, and previous studies have identified male-and hermaphrodite-specific piRNAs by sequencing of whole animals or purified gametes [36,37,52]. We identified 93% of the previously identified male-and 32% of the female-biased piRNAs in another study [52] despite differences between the two (i.e., cell type, genotype, cloning technique and statistics). Thus, our method captures differences in piRNA expression with more sex-specific piRNAs in males than in hermaphrodites, further suggesting sex-specific regulatory mechanisms.

### Endogenous siRNAs with sex-biased germline expression

As a third group of endogenous small RNA, we focused on siRNAs expressed in gonads of both sexes. We analysed all siRNAs that map to protein coding genes with a minimum of five mean antisense reads, thus excluding mRNA degradation products with sense orientation. This revealed 8714 genes with siRNAs in hermaphrodite gonads and 10615 genes in male gonads (Fig 1G). 7675 of these siRNA targeted genes were found in both sexes (Fig 1G; additional file 2). Additional expression data of small RNA (antisense and sense) mapping to other annotated features in the genome such as pseudogenes, transposons and different types of small RNA for gonads of both sexes is available as additional file 2.

To detect potential quantitative expression differences between sexes, we compared normalized antisense read counts per gene between hermaphrodite and male gonads, selecting only genes with at least a four-fold difference in siRNA abundance and a false-discovery rate of 1%. 4508 genes exhibited a sex-biased expression of siRNAs: 1138 were higher expressed in hermaphrodites and 3370 were higher expressed in males. siRNAs of 68 genes were detected exclusively in hermaphrodites and 748 only in males.

To determine whether these expression differences stem from the germline or the somatic gonad, we again made use of the *fog-3* mutants. Of the 4508 genes with sex-biased siRNAs expression in the gonad, 3173 were also differentially expressed between the gonads of control males and feminized males. siRNAs mapping to 2245 genes were more highly expressed in the male gonads, and 928 were more highly expressed in the feminized male gonads (Fig 1H). Thus, most of the sex-differences between hermaphrodite and male gonads stem from expression differences in the germline proper.

To relate sex-differences in germline siRNA expression to biological processes, we carried out gene set enrichment analysis using WormExp v1.0 [53]. This tool queries *C. elegans* mRNA and siRNA expression data in functional groups called ‘gene sets’ based on individual experiments for statistically significant overlap with a given gene list. We predicted that genes with sex-biased germline siRNA expression (i.e., 928 hermaphrodite-and 2245 male-biased genes) may overlap with gene sets corresponding to germline mRNA or siRNA expression. For statistical analysis, we compared enrichment to all 18106 genes with siRNAs detected. From the selected 912 gene sets queried (WormExp categories: mutants, tissue, other), there was significant overlap in both sexes with multiple gene sets related to germline mRNA expression (such as down-regulation in germline-less *glp-1* mutant; here and in the following comparisons p<0.001) and siRNA regulation (such as alteration in *csr-1, rde-1* and *rrf-3* mutants). In addition, this analysis revealed sex-specific overlap with several prior siRNA experiments. Genes with hermaphrodite-biased germline siRNA expression overlapped with siRNA targets detected in whole animals, for example those regulated by *rde-8, ergo-1, eri-6/-7, mut-16*. Likewise, genes with male-biased germline siRNA expression overlapped with spermatogenesis specific *alg-3/-4* targets. Thus, as expected, gonadal siRNAs with tissue-and sex-specific expression are involved in germline and siRNA functions, confirming the specificity of our approach.

### High 22G RNA levels are associated with low mRNA expression, compatible with silencing

Since 22G RNAs of several biogenesis pathways induce gene silencing, we determined the relative abundance of 22G RNAs in the gonads of both hermaphrodites and males. 22G RNAs represented 27.4% of reads in the hermaphrodite gonads and 27.9% of reads in the male gonads (additional file 1 Fig S2A). The 22G RNA levels per gene were not significantly different between the male (mean reads 234.4±1432.9) and hermaphrodite gonads (173.2±857.8; additional file 1 Fig S2B).

To gain insights into 22G RNA-mediated mRNA silencing in gonads, we tested if there was an association of 22G RNAs and corresponding mRNA template expression. If mRNA induces 22G RNA synthesis, there should be a positive correlation between the levels of mRNA and 22G RNA. Alternatively, if 22G RNA expression causes mRNA silencing, a negative correlation between the levels of mRNA and 22G RNA is expected. Data on gonadal mRNA expression in both *C. elegans* sexes was available from a previous study [54]. We first grouped genes in 20 bins according to their 22G RNA expression level. Across bins, there was a positive association between the levels of 22G RNA and gonadal mRNA for groups of genes with low and medium levels of 22G RNA (Fig 2). In contrast, in both hermaphrodites and males, the level of mRNA decreased with extremely high 22G RNA levels (Fig 2), a pattern suggestive of silencing. For statistical analyses we next grouped genes in 5% intervals with equal number of genes again according to 22G RNA expression (additional file 1 Fig S3A). The most significant negative association between the levels of 22G RNA and mRNA was found only for the 5% of the genes with highest level of 22G RNA expression (hermaphrodites p=1×10^-12^, males p=1×10^-16^; additional file 1 Fig S3B). These data indicate that mRNA expression is globally decreased for genes with very high levels of 22G RNAs. In conclusion, the negative correlation of 22G RNA and mRNA expression in *C. elegans* gonads is suggestive of siRNA mediated gene silencing.

**Figure 2:**
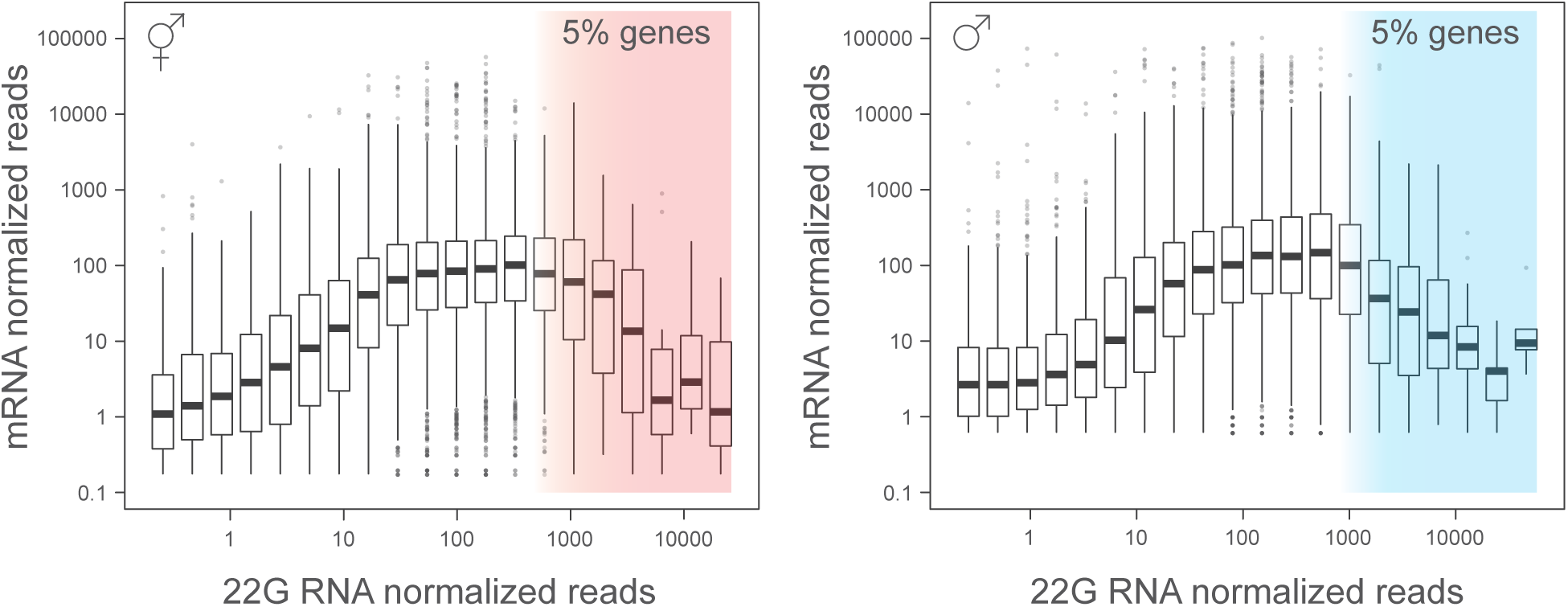
High 22G RNA levels are associated with low mRNA expression, compatible with silencing. Mean normalized mRNA [54] and 22G RNA reads detected in hermaphrodite and male gonads (data in 20 bins according to 22G RNA level; number of genes per bin: 4 – 1101 in hermaphrodites, 5 – 1343 in males; mean: solid line, outliers: open circles). mRNA expression is globally decreased for the top 5% of genes (shaded area) with high levels of 22G RNA. See additional file Fig S3 for statistical analysis.

### Germline sex regulates environmental siRNA accumulation

Since endogenous siRNA pathways share secondary siRNAs owing to similar biogenesis routes [16,18,30], we cannot experimentally determine the fraction of 22G RNA induced by a specific primary siRNA type. To compare expression levels of both primary and secondary siRNA and their impact on protein expression, we developed an sensor based assay. To this end, we used a germline expressed GFP-sensor [55] targeted by *gfp(RNAi)*. Environmental *gfp* dsRNA provided by feeding allows one to manipulate primary siRNA levels and thus directly measure the impact of altered levels of primary siRNA on secondary siRNA levels. To distinguish exogenous primary siRNAs from worm-generated secondary siRNAs, we generated *gfp* dsRNA with single-nucleotide-polymorphisms (SNPs) every 21 nucleotides relative to the GFP-sensor transgene. Thus, primary siRNAs can be discriminated from secondary siRNAs (Fig 3A). To investigate potential sex-specific regulation of siRNA levels during exogenous RNAi, we quantified primary and secondary siRNAs by sequencing siRNAs from gonads of males and hermaphrodites expressing the GFP-sensor. Uptake and primary siRNA processing were active in both male (9 siRNA/ 10^6^ reads) and hermaphrodite gonads (6 siRNA/ 10^6^ reads). Moreover, primary siRNA levels were not statistically different between the sexes (t-test p=0.08, Fig 3B). By contrast, the secondary siRNA level was significantly lower in the male (82 siRNA/10^6^ reads) than hermaphrodite gonads (280 siRNA/10^6^ reads, t-test p=4.3×10^-3^, Fig 3B). Accordingly, the ratio of secondary siRNA/ primary siRNA was significantly lower in the male gonads compared to the hermaphrodite gonads (respectively, 9 and 56 secondary siRNA/primary siRNA, t-test p=0.03, Fig 3B). Taken together, we conclude that environmental supplied dsRNA triggers primary and secondary siRNA production in the germline, with notably higher levels of secondary siRNA products in hermaphrodites than in males. The presence of primary siRNAs in the male germline establish that uptake and transport of silencing agents (such as dsRNA or primary siRNA) from the environment across male somatic tissues, notably the somatic gonad, is functional.

**Figure 3:**
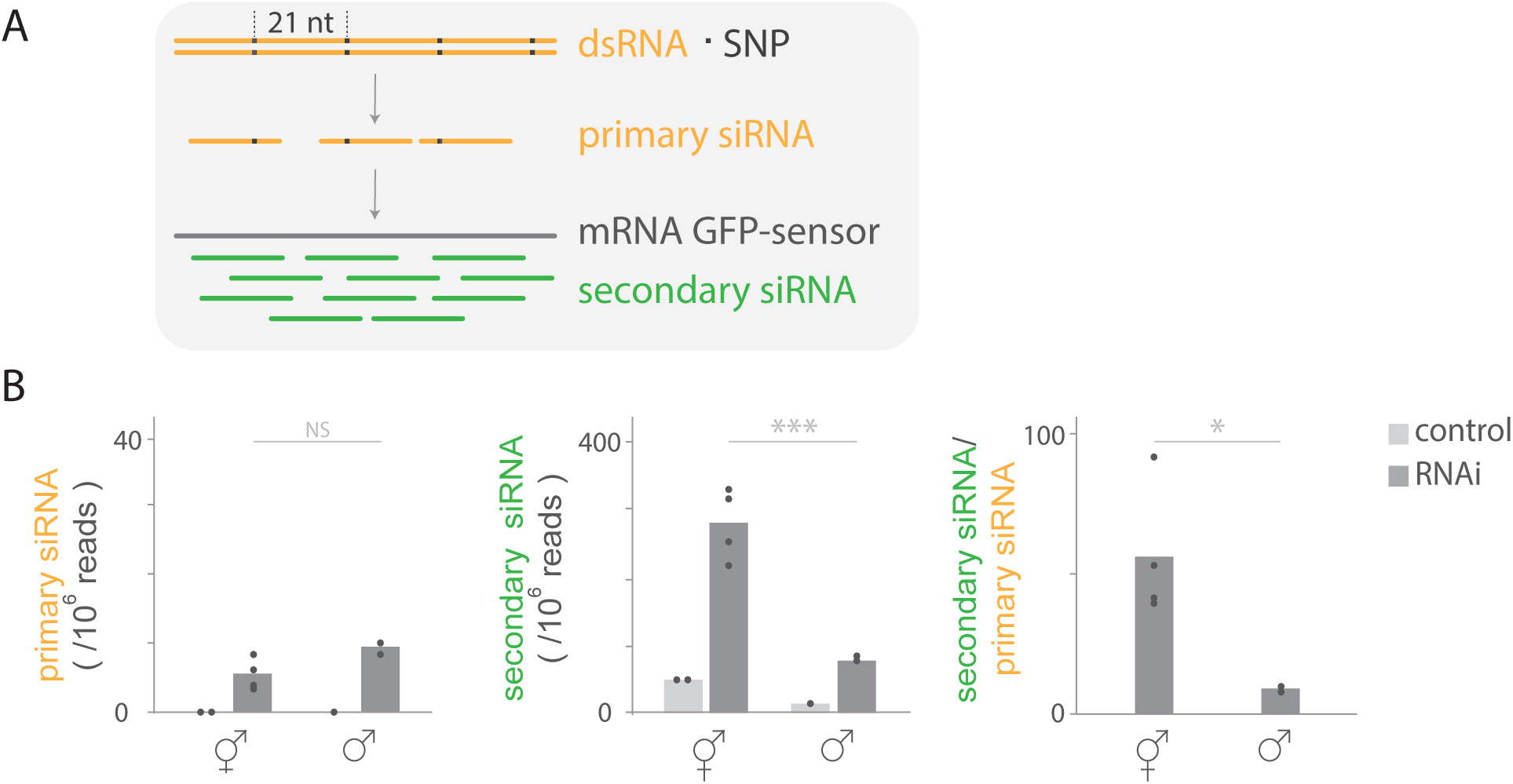
Sex-differences in gonadal siRNA levels during germline RNAi silencing. A) Assay to quantify primary and secondary siRNA levels (from dissected gonads from hermaphrodites and males; see Fig 1B). Animals expressing the GFP-sensor in the germline were raised on *gfp(RNAi)* containing SNPs (black square) each 21 nucleotides. Primary siRNAs (orange) are identified by the presence of the SNPs, while secondary siRNAs (green) lack SNPs. B) Mean primary and secondary siRNA reads normalized to total reads in gonads from hermaphrodites (circles indicate replicates, n=4) and males (n=2). Animals raised on *gfp(RNAi)* or control empty vector (hermaphrodite n=2, male n=1). Mean secondary siRNA/ primary siRNA ratio only shown for RNAi treated animals. T-test was applied for significance test; * = p<0.05, *** = p<0.001.

### RNAi efficacy is germline sex dependent

Anecdotal evidence suggests that germline RNAi silencing may differ between *C. elegans* hermaphrodites and males. RNAi in hermaphrodite germlines is extremely potent as depletion phenotypes often appear within 24 hours of dsRNA exposure [45], even at reduced dsRNA dose [44]. In contrast, similar experiments generally provide negative RNAi results in male germlines [43,46,48]. Presumably stronger RNAi depletion occurs in male germlines directly injected with high doses of dsRNA [56], or by exposing the parental generation [57–60]. However, direct comparison of silencing efficacy of endogenous genes solely based on phenotypes in germlines of hermaphrodites and males is insufficient, notably due to sex differences in physiology and gene function. Feeding dsRNA triggers against the expressed GFP-sensor allows one to measure and compare siRNA levels and resulting RNAi silencing in hermaphrodites and males raised in the same environment. The observed difference in siRNA amplification products observed above (see Fig 3) prompted us to ask whether germline sex affects RNAi silencing. To assess RNAi silencing in both sexes, we monitored the presence or absence of GFP-sensor fluorescence upon *gfp(RNAi)* (Fig 4A). These analyses revealed that germline silencing by dsRNA is sex specific. Of the 307 hermaphrodites analysed, 305 (99.4%) silenced the germline GFP-sensor (Fig 4B), in contrast to only 3 (3.7%) of the 81 males analysed (chi-square p<0.001; Fig 4B). This difference pertained to the entire germline of males since GFP-sensor expression was visible in proliferating germ cells and differentiated spermatocytes (Fig 4A). Contrary to the germline, the soma of males was RNAi sensitive, as evidenced by the complete silencing of a ubiquitous GFP-sensor [61] in all non-neuronal somatic cells (Fig 4C; 60/60 males). This is in line with successful RNAi-based screens targeting somatic tissues in male, notably the somatic gonad [62,63] and shows that somatic RNAi efficacy is sex independent. Thus, isogenic worms raised in the same environment show phenotypic differences in response to environmental cues. Since RNAi silencing is dependent on the dsRNA levels and because mutants defective in siRNA amplification are partly RNAi resistant [16,64–66], the observed sex-difference in secondary siRNA abundance in gonads (Fig 3B) may explain the difference between sexes in germline RNAi efficacy (Fig 4A and 4B).

**Figure 4:**
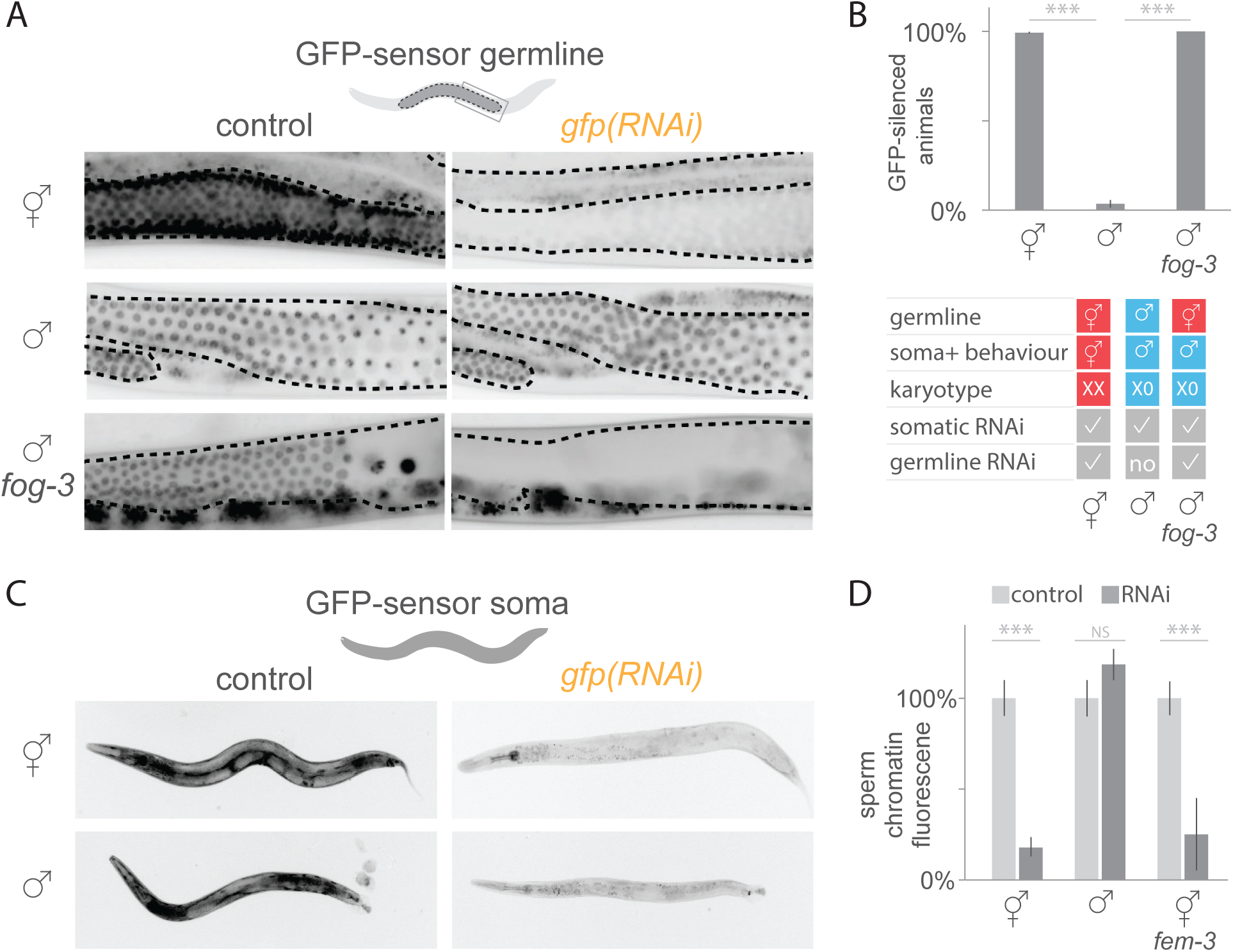
Sex-differences in germline RNAi efficacy. A) Fluorescence microscopy images of germline (dotted line) GFP-sensor expression in hermaphrodites, males and *fog-3* males (left). Silencing of the GFP-sensor upon *gfp(RNAi)* in hermaphrodite and *fog-3* male germlines, while control male germlines are RNAi resistant (right). Gut autofluorescence is visible outside the germline. B) Percentage of worms (±SEM) silencing the GFP-sensor in the germline upon *gfp(RNAi)* (n=307 hermaphrodites, 81 males, 106 *fog-3* males). Chi-square test was applied; *** = p<0.001. Table with hermaphrodite/ female (red) and male (blue) traits in hermaphrodites, males and *fog-3* males. C) Fluorescence microscopy images of ubiquitous GFP-sensor expression in the soma of hermaphrodites and males (left, for both n=54). *gfp(RNAi)* silencing of somatic GFP in non-neuronal cells was observed in all hermaphrodites and males (right, respectively n=57 and n= 60). D) Background subtracted sperm GFP-sensor chromatin signal intensity (mean ±SE) upon *gfp(RNAi)* in hermaphrodites (n=10), males (n=9) and *fem-3* hermaphrodites producing only sperm (n=15). Signal intensity was normalized to animals not exposed to dsRNA (n=10 hermaphrodites, 11 males, 12 *fem-3* hermaphrodites). T-test was applied for significance test; *** = p<0.001.

### RNAi efficacy is germline sex dependent

To determine whether the observed differences in germline RNAi silencing between male and hermaphrodites was induced by differences in the germline or the soma, we again made use of the feminized males. All other traits of the feminized males such as soma, karyotype and feeding-behaviour are undistinguishable from wild-type males (Fig 4B). Upon feeding *gfp(RNAi)*, all 106 feminized males silenced the GFP-sensor (Fig 4A, B), indicating that germline-intrinsic factors regulate RNAi silencing, rather than other male traits such as the soma, karyotype or behaviour.

To test whether differences in germline RNAi silencing between control and feminized mutant male gonads could stem from altered siRNA levels or altered downstream silencing activities, for instance target slicing by Argonautes, we compared siRNA levels in gonads. The gonads of feminized males contained higher levels of primary *gfp* siRNAs (74 siRNA/10^6^ reads, t-test p=9.5×10^-4^) and secondary *gfp* siRNAs (798 siRNA/10^6^ reads, t-test p=0.04) than control male gonads (additional file Figure S4). Since RNAi silencing is dose-dependent, the higher siRNA levels in feminized mutant compared to male gonads are a plausible explanation for the sex-difference in germline RNAi silencing.

To examine whether the male germline RNAi resistance was inherent to sperm or influenced by the surrounding germline environment, we compared RNAi silencing between the sperm of males and hermaphrodites. If the germline environment regulates RNAi, silencing should differ between the sperm of males and hermaphrodites. By contrast, if the sperm physiology confers RNAi resistance, both types of sperm should be RNAi resistant. We measured fluorescence intensities of a GFP-sensor [67] on chromatin using both types of sperm imaged in whole worms fed with *gfp(RNAi)*. This experiment revealed RNAi silencing in hermaphrodite sperm, since *gfp(RNAi)* significantly decreased sperm GFP signal intensity by a mean of 81.9%±5.1% compared to animals not exposed to dsRNA (Fig 4D; t-test p=1.8×10^-6^). By contrast, the sperm of males was completely RNAi resistant, with the GFP-sensor signal intensity being not significantly different between males fed with *gfp(RNAi)* and males not being exposed to dsRNA (Fig 4D; t-test p=0.2).

The finding that, in contrast to sperm in males, sperm from hermaphrodites is RNAi sensitive, indicates that the surrounding germline environment plays a crucial role in RNAi silencing of sperm. Because germ cell precursors for sperm and oocytes grow in the same syncytium in hermaphrodites, silencing agents such as siRNAs or RNAi proteins produced in oocytes may diffuse in the germline and thus influence silencing in sperm. To test this hypothesis, we investigated whether mutant hermaphrodites producing only sperm initiate RNAi independently of oogenic germ cells using the gain-of-function allele *fem-3(q96)* [68]. We found that such sperm were RNAi sensitive, since fluorescence GFP-sensor [67] signal intensity was on average 74.8%±19.9% lower in *fem-3* worms fed *gfp(RNAi)* compared to animals not exposed to RNAi (Fig 4D; t-test p=3.0×10^-7^). Thus, the sperm of hermaphrodites is RNAi sensitive even in the complete absence of oogenic germ cells in the syncytium.

## DISCUSSION

Here we provide the first comprehensive small RNA profile of isolated *C. elegans* male and hermaphrodite gonads and show qualitative as well as quantitative sex-differences in siRNA expression for both somatic gonad and germline tissue. We demonstrate tissue-specific sex-differences in response to environmental RNAi triggers: in contrast to the hermaphrodite germline, the male germline is resistant to silencing and accumulates lower levels of RNAi amplification products. Taken together, these results provide a mechanistic explanation for sex-differences in RNAi efficacy in response to the environment with implications for trans-generational inheritance.

### Sex-differences in gonadal miRNA, piRNA and siRNA expression

miRNA regulation, notably via the conserved miR-35 family is crucial for the embryonic viability, proliferation of the germline and is implicated in sex-determination [27,28,41,69]. In addition, miRNAs expressed in the *C. elegans* somatic gonad maintain germline proliferation and differentiation [41]. We identified female germline-biased expression of the miR35-41 cluster in isolated gonads, which is in line with studies conducted on whole animals [27,29,70].

The transgene reporter for miR-246 expression, which typically escapes germline expression, was previously observed in hermaphrodite gonadal sheath cells [23]. Here we establish sex-biased expression of endogenous miR-246 specifically in the hermaphrodite somatic gonad as opposed to the germline or the male gonad. Since miR-246 expression positively correlates with lifespan [71,72], it will be interesting to identify miR-246 targets in the somatic gonad or germline. Thus, small RNA sequencing of isolated gonads and comparison of expression differences between wild-type and sex-transformed mutant gonads captures known sex-differences in expression and in addition provides more detailed, tissue-specific information.

Our data provide the first tissue-specific analysis of miRNA expression in males. Of the 46 male-biased miRNAs with searchable names in miRBase detected in whole animals [24], 13 miRNAs showed male-biased gonad expression, underlining the importance of a tissue-specific approach. Thus, sex-biased miRNA expression occurs in both soma and gonad. Some miRNAs with male-biased gonad expression were previously not detected as sex-biased [24] and could have been masked by somatic expression in both sexes.

In contrast to miRNAs, the piRNA pathway acts exclusively in the germline. Since the germline makes up about half of the adult *C. elegans* cells [73], whole worm sequencing is a good proxy for germline-restricted piRNA expression. Our analysis of sex-biased piRNA expression thus complements previous studies on whole animals that reported distinct piRNA expression in hermaphrodites and males [24,37]. While the cloning method applied here causes relatively low (5’-monophasphate) piRNA detection level compared to whole animals [37], it also causes preferential cloning of 5’-triphosphate RNA species and thus high coverage of endogenous siRNA. The tissue-specific analysis of endogenous 22G RNA and mRNA identified transcripts that are potentially regulated by RNAi. Importantly, hermaphrodites and males share common as well as distinct siRNA targeted gene sets in gonadal tissue. Most gonadal sex-differences in siRNA expression stem from the germline, which represents the vast majority of the gonadal tissue. Further comparison of gonadal siRNA and corresponding mRNA uncovers uncoupling of siRNA and mRNA levels, indicative of RNAi silencing. It will be interesting to investigate this regulation in the future.

### RNAi efficacy is regulated by germline sex

Our study provides novel insights into how the sex of an animal influences RNA-induced environmental changes in the phenotype. In particular, using environmental dsRNA triggers, we demonstrated that RNAi resistance affects exclusively the male germline. Sequencing primary and secondary siRNA with unique SNPs in isolated gonads from males and hermaphrodites allowed us to quantitatively compare steady-state levels. We detected similar levels of primary siRNAs in gonads of both sexes, suggesting functional transport of RNAi trigger from the environment to the soma and processing of primary siRNA in the male germline. In contrast, we discovered low levels of secondary siRNA amplification products in males compared to hermaphrodites. Thus, RNAi resistance in the male germline affects downstream processes that likely impact sex-differences in secondary siRNA processing or siRNA stability. It follows that RNAi is generally not suitable for functional studies of the *C. elegans* male germline if provided during one generation only. Since RNAi silencing is efficient in sperm of hermaphrodites, such animals are more suitable for reverse genetics targeting the germline.

What determines the sex-differences of gene regulation in response to environmental cues? It is conceivable that known or novel RNAi regulators impact germline gene regulation by modulating RNAi efficacy. A potential candidate is the endoribonuclease RDE-8 which is expressed predominantly in hermaphrodite germlines and is essential for amplification of siRNA products and silencing [66]. Thus, RNAi resistance in male germlines may possibly be linked to reduced RDE-8 activity. Also, sex-differences in localization may affect RNAi efficacy. Notably CID-1, a poly(U)-polymerase modulating germline RNAi efficacy shows distinct subcellular localization in male and female germlines – perinuclear, respectively chromosomal-that may affect function [74]. Intriguingly, in addition, multiple sex-differences are well described for the Argonaute CSR-1 that targets endogenous germline transcripts via secondary 22G RNAs [19,34]. First, males and hermaphrodites express distinct CSR-1 isoforms [34,73]. Second, *csr-1* mutant hermaphrodites are mostly sterile, while *csr-1* mutant males show only modestly reduced fertility over multiple generations [33,75]. Third, CSR-1 bound 22G RNAs target distinct sets of transcripts in male and hermaphrodite [33,75]. Thus, sex-differences in CSR-1 function may contribute to the observed sex-difference in RNAi sensitivity, in line with CSR-1 function in germline RNAi [76]. Overall, multiple mechanisms may contribute to sex-specific gene regulation and untangling the individual contributions will provide exiting answers on male and female biology.

In conclusion, this study provides the tissue-specific small RNAome of *C. elegans* hermaphrodite and male gonads and identifies quantitative sex-differences in miRNA, piRNA and siRNA expression. Furthermore, we demonstrate that the male germline is resistant to RNAi triggers taken up from the environment and accumulates lower levels of RNAi amplification products. We thus provide mechanistic insights into sex-differences of gene regulation in response to environmental cues that may play a role in transgenerational inheritance.

## MATERIAL AND METHODS

### Nematode strains

*C. elegans* strains of the following genotype were cultured according to standard procedures [77]: *him-8(e1489)* IV [77], *fog-3(q849[E126K])* I/ hT2*[qIs48]* (I;III); (kind gift from Scott Aoki and Judith Kimble) and *fem-3(q96)* IV [68]. The following GFP-sensors were used: single copy *mjIs145* [mex-5p::GFP::his-58::21UR-1sense::tbb-2 3′UTR] II (Bagijn 2012; Figure 1, 2, 3, 4A, B), multicopy *adIs2122* [lgg-1p::GFP::lgg-1 *3’UTR* + rol-6(su1006)] [61]; Figure 4C) and multicopy *zuIs178* [*his-72p::his-72::GFP:his-72 3’UTR*] ([67]; Figure 4D). All strains were wild-type for *mut-16(mg461)* by PCR [78]. *him-8(e1489)* was used to generate otherwise wild-type males and hermaphrodites, for simplicity we refer to such animals as ‘control males’ and ‘control hermaphrodites’.

### RNA interference

RNAi feeding plates where prepared from freshly streaked HT115 bacteria containing L4440 vector according to [79]. Those were used to feed synchronized L1 worms minimum 48h at 20°C unless otherwise stated. Initial *gfp(RNAi)* experiments were carried out with a construct targeting the full *gfp* coding sequence including three introns. The recoded *gfp(RNAi)* with SNP every 21 nucleotides targets only exons and was synthesized by Integrated DNA Technologies and cloned into the L4440 plasmid. Silencing efficiency of recoded *gfp(RNAi)* is slightly reduced compared to full length *gfp(RNAi)*, which may be caused by SNPs or lacking exons.

### Microscopy

RNAi silencing was scored in live adults mounted on slides with reaction wells (Paul Marienfeld) using 20x or 40x objective on a Zeiss Axio Scope.A1 microscope. Worms scored as RNAi sensitive on a binary scale if GFP-sensor was absent or greatly diminished as compared to animals treated with empty vector RNAi. Differential interference contrast (DIC) and fluorescence images were acquired on a Axiocam 506 mono CCD camera and processed with Fiji software [80]. Outlines of the germlines where drawn on the DIC image in Adobe Illustrator.

### Gonad isolation and replicates

Bleach-synchronized L1 worms were grown on RNAi bacteria 48h at 20°C until L4/ young adult, then washed 5 times in 15mL M9 to remove bacteria and grown overnight at 20°C on OP50 bacteria. Prior dissection, adults were picked on empty plates and transferred in groups of 3-4 in a drop of 0.01% levamisole + sperm buffer [81] to wash and paralyze animals. Animals were cut with a 21G needle behind the pharynx to liberate the gonad in a new drop of buffer on a depression slide (cavity 15-18mm, depth 0.6-0.8 mm; Marienfeld). Non-gonadal somatic tissue was removed, including the intestine; the spermatheca was removed from hermaphrodite gonads. Using a mouth pipette with glass capillary gonads where transferred to a tube with RNAlater (50-100µl) and after TRIzol (Thermo Fisher) addition frozen at −80°C.

Gonads from hermphrodites (100), males or *fog-3* males (each 200) were processed as independent replicates for sequencing as follows (# replicates; including # recoded *gfp(RNAi)* samples): hermaphrodites (6;4), males (3;2) and *fog-3* males (2;2). Since endogenous siRNAs levels were not different from germlines treated with or without *gfp(RNAi)*, both were used for sex-biased expression analysis.

### RNA extraction

RNA was extracted with 1 ml of TRIzol according to manufactor’s protocol. Briefly, RNA was precipitated adding 1 volume of isopropanol and 20 µg glycogen (Roche). Samples forming a white precipitate at this point were cleared by addition of 500 µL isopropanol: water 50 % (v/v). Samples were frozen at −80°C, thawed on ice and RNA was pelleted by centrifugation for 30 min at +4°C, 16000 g. The pellet was washed in ice cold 70% (v/v) ethanol and recovered by centrifugation for 20 min at +4°C, 16000 g and finally resuspended in 10 µL water. RNA concentration was determined by Qubit® RNA BR Assay Kit (Thermo Fisher).

### 5’ independent library preparation and sequencing

For RNA dephosphorylation 400 ng of RNA were treated with 20 Units of 5’ polyphosphatase (Epicenter) in 20 µL reaction volume, purified using acid-phenol-chloroform, pH 4.5 (Thermo Fisher) and isopropanol precipitated using 20 µg of glycogen. Subsequently, RNA was suspended in 6 µL water and directly used for TruSeq Small RNA library kit (Illumina) following the manufacturer’s instructions with exception that 15 cycles of PCR amplification were used. The cDNA libraries were separated on 6% TBE PAGE gels (Life Technologies) and bands with 147-157 nucleotides were cut from the gel. The gel matrix was broken by centrifugation through gel breaker tubes (IST Engineering Inc.) and size-selected cDNA eluted with 400 μl of 0.3M Na-Chloride. cDNA was purified by centrifugation through Spin-X 0.22μm cellulose acetate filter columns (Costar) followed by isopropanol precipitation. Libraries were sequenced on a HiSeq 1500 Sequencer (Illumina).

### Computational analysis of small RNA high-throughput sequencing data

Small RNA sequencing results were obtained from https://basespace.illumina.com/ as fastq files after demultiplexing. Sequencing data is available in the European Nucleotide Archive under accession number PRJEB12010. Firstly, adapter sequences, reads shorten than 21 nucleotides (nt) and reads longer than 34 nt were removed using cutadapt v1.9.1. Secondly, the remaining reads were aligned using bowtie 1.1.2 with at most 2 mismatches to *Escherichia coli* str. K-12 MG1655. Thirdly, the remaining reads were aligned with at most 2 mismatches to *C. elegans* WS235 tRNAs and rRNAs. Fourthly, the remaining reads were aligned with no mismatches to the worm *gfp* sequence. Fifthly, the remaining reads were aligned with no mismatches to the recoded *gfp* sequence. Sixthly, the remaining reads were aligned with no mismatches to *C. elegans* miRNAshairpins (miR-base release 21). Seventhly, the remaining reads were aligned with no mismatches to *C. elegans* genome cel235. Overall, we obtained ∼2.3 – 12.8 million trimmed reads mapping to the genome (WS235) for each sex. endo-siRNAs was quantified per gene, antisense reads mapping to coding exons, which account for more than 96% of reads in this class.

#### Silencing of mRNA targeted by very high levels of 22G RNA

Fastq-mcf (Erik Aronesty (2011). ea-utils : ″Command-line tools for processing biological sequencing data″; https://github.com/ExpressionAnalysis/ea-utils) was used to trim adapters from single-end reads. Bowtie 1.1.2 (parameters “--best --strata --tryhard -m 1”) (Langmead B, Trapnell C, Pop M, Salzberg SL. <u>Ultrafast and memory-efficient alignment of short DNA sequences to the human genome</u>. *<u>Genome</u> <u>Biol</u>* 10:R25.) was used to map single-ended reads to *C. elegans* WS250 genome. Reads were classified as 22G/26G based on length and starting base. HTSeq 0.6.0 (Simon Anders, Paul Theodor Pyl, Wolfgang Huber *HTSeq — A Python framework to work with high-throughput sequencing data* Bioinformatics 2014) was used to generate counts for reads that map to WS250 features in the sense and antisense directions. Reads mapping against transposons were retrieved using c_elegans.PRJNA13758.WS250.annotations.exons.genes.with_transposons. gff. Features counts from separate samples were normalized and differences in expression were determined using DESeq2 1.13.8 (Love MI, Huber W and Anders S (2014). “Moderated estimation of fold change and dispersion for RNA-seq data with DESeq2). Because alternative biological processes generate sense and antisense mapping reads, reads mapping in the sense direction from each library were normalized together, likewise reads mapping in the antisense direction were separately normalized. The cutoff for sex-biased expression was >4 fold difference in abundance and adjusted p-value <0.01. For correlations with germline transcriptome, data was used from West et al., Genome Biology 2017 (Sequence Read Archive, accession number SRP096640). For miRNAs both star and non-star sequences from the same gene were summed.

#### Silencing of mRNA targeted by very high levels of 22G RNA

Assay and quantification of primary and secondary *gfp* siRNA levels: Reads were trimmed with cutadapt v1.9.1. Bacteria, tRNA and rRNA reads contaminations were removed using bowtie 1.1.2 alignment to the *Escherichia coli* str. K-12 MG1655 and to *C. elegans* WS235 tRNAs and rRNAs. Remaining reads were aligned with no mismatches to the worm *gfp* sequence. Then the remaining reads were aligned with no mismatches to the recoded *gfp* sequence.

## LIST OF ABBREVIATIONS

(RNAi): RNA interference
(nt): Nucleotide
(RISC): RNA-induced silencing complex
(RdRP): RNA-dependent RNA polymerases
(piRNA): PIWI bound small RNA
(gfp): Green fluorescent protein
(SNP): Single-nucleotide-polymorphism

## DECLARATIONS

### Ethics approval and consent to participate

Not applicable.

### Consent for publication

Not applicable.

### Availability of data and material

Small RNA library sequencing data is available on the European Nucleotide Archive in the Study: PRJEB12010. A sample description can be found in additional file 3.

### Authors’ contributions

AB and FB contributed equally to experimental design, collected data, analysed and interpreted results, and drafted the manuscript. SW contributed to analysis design and interpretation of data and results. RC and AD contributed to cloning and data collection with GFP-sensor strains. FS provided statistical analysis for Fig 2. PG, KG and FP were involved in the conception and design of experiments and analyses. LK and EAM contributed to design of experiments and analyses, results interpretation and drafting the manuscript. All authors read and approved the final manuscript

## Acknowledgements

We are grateful to Scott Aoki and Judith Kimble for sharing strains prior to publication, Benita Wolf, Adria LeBoeuf and Tamara Mikeladze-Dvali for comments on the manuscript and Gönczy lab members for advice. We thank Kay Harnish of the Gurdon Institute Sequencing Facility for managing the high-throughput sequencing, Nazife Bega for media preparation and Wormbase. Some strains were provided by the CGC, which is funded by NIH Office of Research Infrastructure Programs (P40 OD010440).

## ADDITIONAL DATA

### Additional file 1.pdf

Supplemental figures S1-S4.

### SUPPLEMENTAL FIGURE LEGENDS

**Figure S1:**
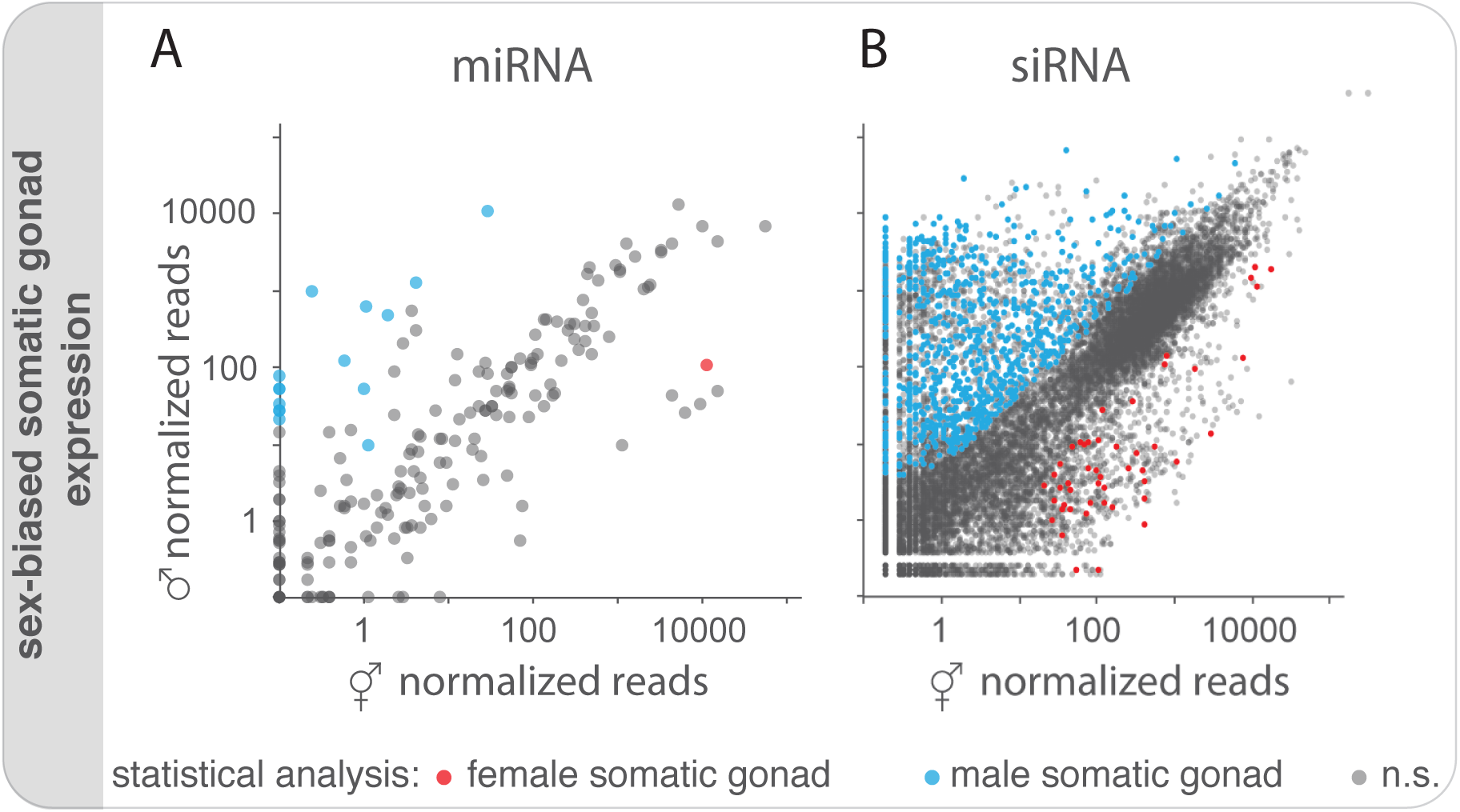
Somatic gonad sex-differences in expression of miRNA and endogenous siRNA. A) Mean normalized miRNA reads (sense) in hermaphrodite (n=6 replicates) and male gonads (n=3). miRNA expression-differences with four-fold difference in abundance that were statistically different (Wilcoxon rank sum test with continuity correction; p adjusted<0.01) between gonads of hermaphrodites and males as well as between hermaphrodites and *fog-3* males (n=2) are highlighted in red (female somatic gonad-biased) and blue (male somatic gonad-biased). B) Mean normalized endogenous siRNA reads (antisense) in hermaphrodite and male gonads; same analyses and representation used as in Fig S1A.

**Figure S2:**
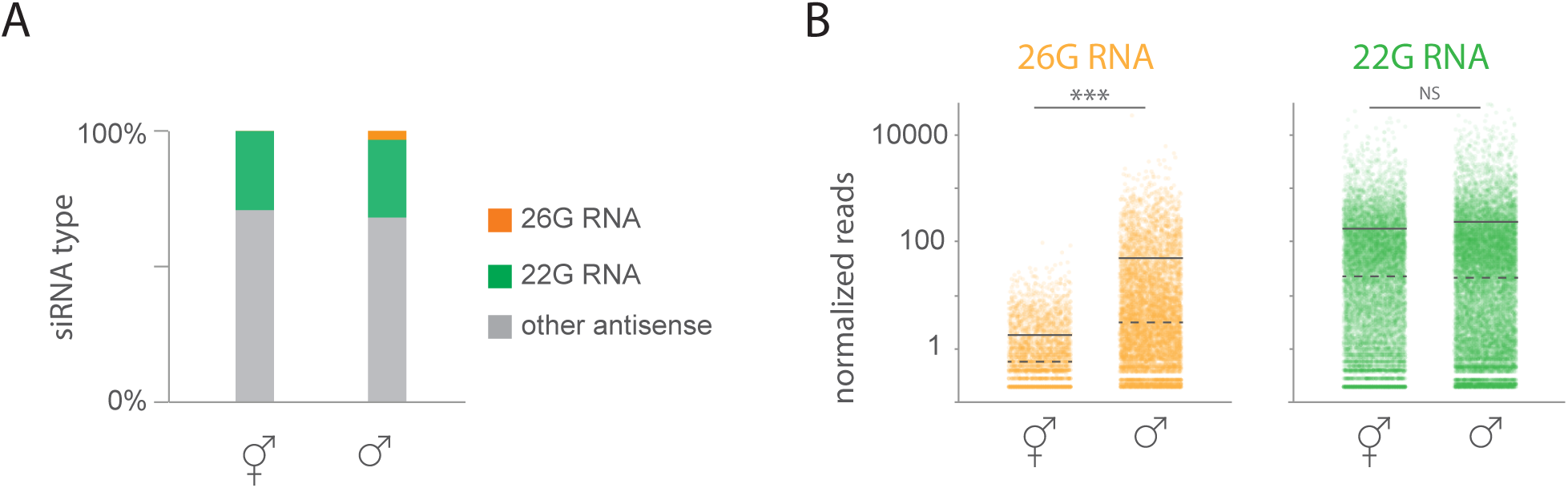
Hermaphrodite and male gonadal 26G RNA and 22G RNA. A) Percentage of antisense reads in hermaphrodite and male gonads classified as primary 26G RNA, secondary 22G RNA or other siRNAs type. B) Normalized mean 26G RNA and 22G RNA reads in hermaphrodite and male gonads per gene.

**Figure S3:**
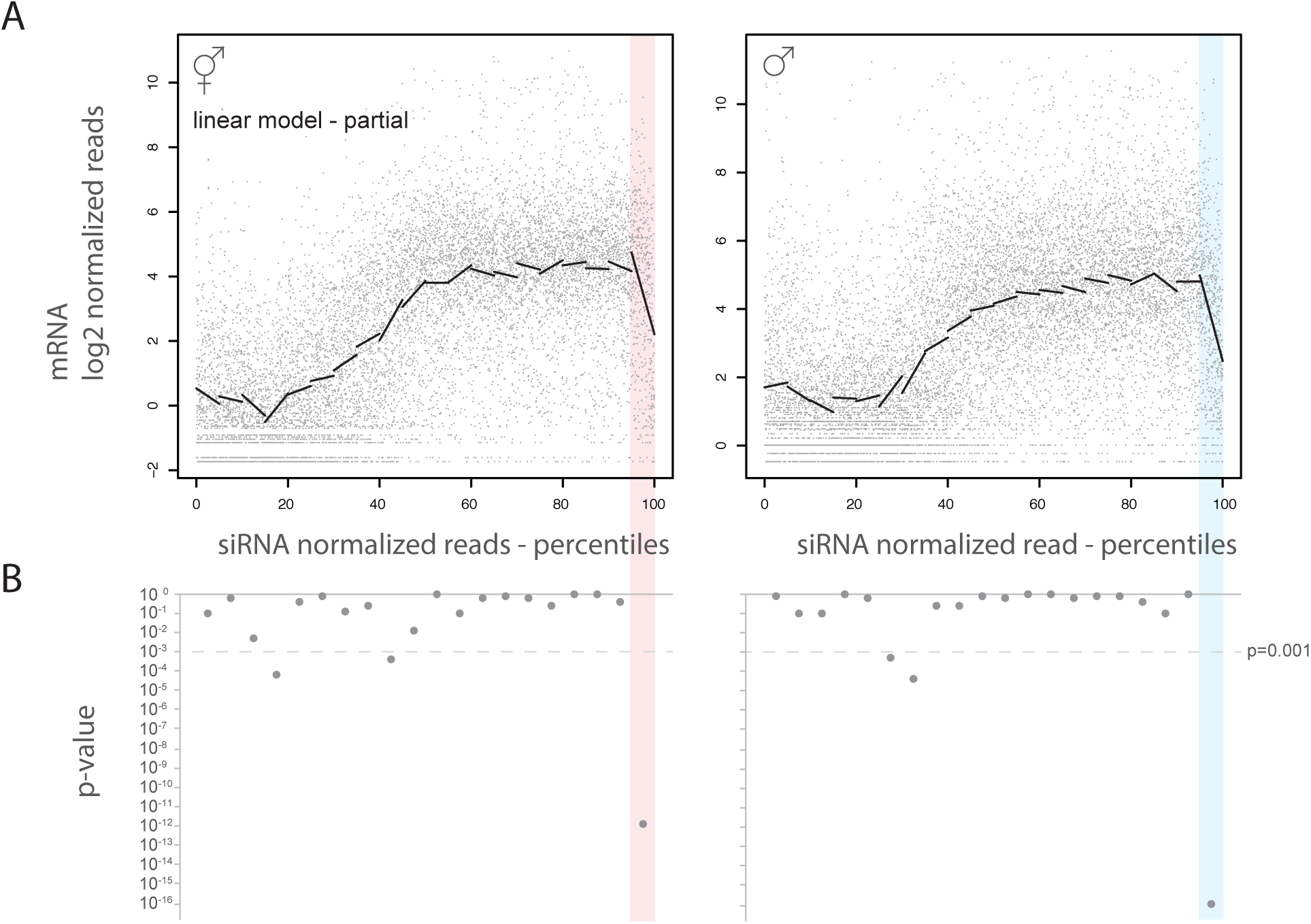
Statistical analysis of gonadal mRNA and siRNA expression levels. A) Mean normalized mRNA reads [54] and siRNAs in percentiles in hermaphrodite and male gonads. For 20 bins (each representing 5% of data i.e. 427 genes per bin in hermaphrodites and 504 in males) a linear model was fitted (black line) B) p-values indicate whether the linear model deviates from zero. The most significant negative correlation between the levels of 22G RNA and mRNA was found for the 5% of the genes with highest level of 22G RNA expression (shaded).

**Figure S4:**
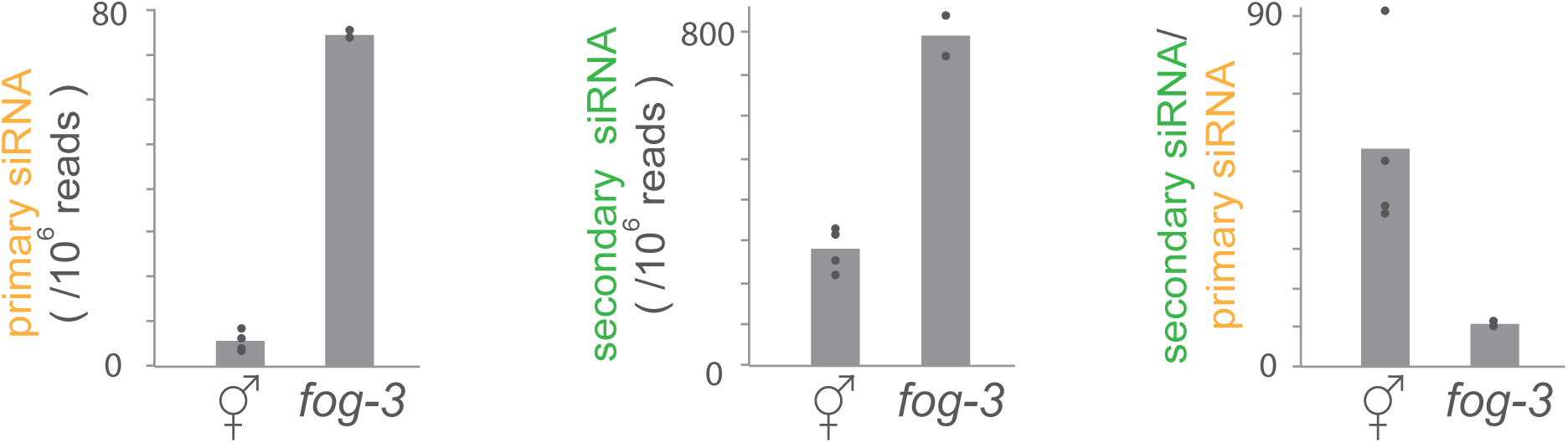
Sex-differences in RNAi efficacy in female germlines. Mean primary and secondary siRNA reads normalized to total reads in gonads from hermaphrodites (circles indicate replicates n=4; data from figure 3B) and *fog-3* males (n=2) raised on *gfp(RNAi)*. Mean secondary siRNA/ primary siRNA ratio. T-test was applied for significance test; * = p<0.05, *** = p<0.001.

**Additional file 2.xls:** Summary gonadal small RNA expression.

Analyses of sex-biased miRNA, piRNA, germline siRNA, sense and antisense reads

**Additional file 3.xls:** Sequencing details.

Accession and genotype of each replicate

